# The effects of biological sex on estimates of persistent inward currents in the human lower limb

**DOI:** 10.1101/2022.10.09.511486

**Authors:** Sophia T. Jenz, James A. Beauchamp, Matheus M. Gomes, Francesco Negro, CJ Heckman, Gregory E.P. Pearcey

## Abstract

Non-invasive recordings of motor unit (MU) spike trains help us understand how the nervous system controls movement and how it adapts to various physiological conditions. The majority of study participants in human and non-human animal physiology studies are male, and it is assumed mechanisms uncovered in these studies are shared between males and females. However, sex differences in neurological impairment and physical performance warrant the study of sex as a biological variable in human physiology and performance. To begin addressing this gap in the study of biophysical properties of human motoneurons, we quantified MU discharge rates and estimates of persistent inward current (PIC) magnitude in both sexes by quantifying ΔF. We decomposed MU spike trains from the tibialis anterior (TA), medial gastrocnemius (MG), and soleus (SOL) using high-density surface electromyography and blind source separation algorithms. Ten participants of each sex performed slow triangular (10s up and down) isometric contractions to a peak of 30% of their maximum voluntary contraction. We then used linear mixed effects models to determine if peak discharge rate and ΔF were predicted by the fixed effects of sex, muscle, and their interaction. Despite a lack of significant sex-differences in peak discharge rates across all muscles, ΔF was larger (χ^2^_(1)_ = 6.26, *p* = 0.012) in females (4.73 ± 0.242 pps) than males (3.81 ± 0.240 pps). These findings suggest that neuromodulatory drive, inhibitory input, and/or biophysical properties of motoneurons differ between the sexes and may contribute to differences in MU discharge patterns.

**KEY POINTS:**

– *Sex differences in motor unit studies have been revealed with greater inclusion of female participants, however, mechanisms for these differences remain unclear*.
– *Estimates of persistent inward currents (i.e*., ΔF) *were greater in females than males in the tibialis anterior, medial gastrocnemius, and soleus muscles.*
– *This suggests that neuromodulatory drive, monoaminergic signaling, or descending control may differ between the sexes.*
– *Therefore, sex differences in estimates of PICs may provide a mechanism behind previously reported sex differences in motoneuron discharge patterns.*.

## INTRODUCTION

Historically, studies in physiology, and in particular neurophysiology, have excluded female participants. Woitowich and colleagues (2020) recently surveyed the literature and showed that females currently represent only 15 percent of both non-human subjects and human participants in biological studies. This has resulted in an extensive understanding of the neural control of movement in males, but a poor understanding of what mechanisms underlie differences in motor control observed in females (Jenz & Pearcey, 2022).

Regardless of biological sex, all movement is achieved through the activation of motoneurons and the muscle fibers they innervate, together known as motor units (MUs; Heckman & Enoka, 2012). Under normal conditions, motoneurons have a one-to-one spike ratio with the fibers they innervate, and therefore the discharge patterns of motoneurons and their MU are the same (Johnson *et al*., 2017; Thompson *et al*., 2018). This creates a unique opportunity to study the central nervous system, as motoneurons are the only neuron that can be recorded non-invasively in humans. Thus, the non-invasive assessment of MU behavior can reveal important information on how the cortex, spinal cord, and individual neurons communicate and whether sexual dimorphisms exist within these neural commands.

A contemporary understanding of the physiology underlying motor control suggests that motoneurons receive a complex combination of excitatory, inhibitory, and neuromodulatory input (Heckman et al. 2009). This neuromodulatory input involves two important monoamines (among others), serotonin (5HT) and norepinephrine (NE), that are released via long descending projections from the brainstem nuclei (raphe nucleus and locus coeruleus, respectively) onto motoneurons (Montague *et al*., 2013). Release of 5HT and NE facilitates voltage-sensitive persistent inward currents (PICs; Hounsgaard *et al*., 1988; Heckman *et al*., 2005) and contributes to the gain control required for humans to perform an impressive variety of movements, from long term postural control during waking for hours, to precise targeted movements like reaching to grasp a pint in a crowded pub (Heckman *et al*., 2008).

PICs modulate motoneuron discharge in a non-linear fashion by creating a strong initial acceleration and subsequent attenuation in discharge rate during linear increases in synaptic input (Binder *et al*., 2020). The most robust hallmark of PICs in humans, however, manifests as discharge rate hysteresis between the recruitment and de-recruitment of MUs (Khurram *et al*., 2022). Typically, PICs are estimated in humans using a paired MU analysis technique to quantify discharge rate hysteresis (Gorassini *et al*., 1998, 2002*a*, 2002*b*). Although many labs have utilized this technique with MU spike trains obtained while participants perform voluntary triangular shaped contractions, the role of descending monoaminergic (i.e., 5HT and NE) input from the brainstem is broadly understudied in the motor system and has not been compared between the sexes.

In recent years, studies that have included an adequate number of females have demonstrated sex differences in several MU discharge properties, but the results vary across different tasks and conditions. The consensus is that MU discharge (English & Widmer, 2003; Harwood *et al*., 2014; Peng *et al*., 2018; Parra *et al*., 2020; Inglis & Gabriel, 2020; Nishikawa *et al*., 2022) and neuromuscular fatigue (Hunter & Enoka, 2001; Celichowski & Drzymała, 2006; Ansdell *et al*., 2019; Inglis & Gabriel, 2021; Augsburger *et al*., 2022) differ between the sexes (Lulic-Kuryllo & Inglis, 2022). Recent data suggest that females use different recruitment strategies than males at relatively low efforts due to smaller MU size and higher discharge rates (Guo *et al*., 2022) but the specific mechanisms underlying these differences remain unknown.

In non-human animal models, sex differences in MU physiology have been identified at the cellular level. For example, male rats have more fast-fatigable type MUs in the gastrocnemius muscle, while female rats have more fatigue-resistant MUs (Celichowski & Drzymała, 2006), with male rats having a greater total number of MUs than females (Celichowski & Drzymała-Celichowska, 2007). Most relevant to the current investigation, Kirkwood and colleagues (Kirkwood *et al*., 2002; Ford & Kirkwood, 2006) have shown that plateau potentials, which are dependent on Ca^2+^ PICs and typically absent in anesthetized preparations, have been observed only in anesthetized estrous *female* cats, suggesting that biophysical motoneuronal properties differ between the sexes.

Therefore, to understand the effect of biological sex on motoneuronal properties in humans, we measured MU behavior during slow isometric ramps contractions to the same relative intensity and compared them between females and males. We hypothesized that females would have greater PICs, which would manifest as higher peak discharge rates and more discharge rate hysteresis – an indication of greater PIC magnitudes.

## METHODS

### Participants and ethical approval

Young and healthy females (n = 10; 25.5 ± 4.8 years) and males (n = 10; 27.2 ± 5.8 years) were recruited for this study. Participants had no history of muscular or neurological impairment and provided written informed consent, approved by the Institutional Review Board of Northwestern University (STU00216360).

### Experimental Apparatus

#### Torque

During each session, participants were seated in a Biodex chair (Biodex Medical Systems, Shirley, NY) and their right foot was placed in a footplate attachment with an ankle angle of 95 degrees, a hip flexion angle of 100 degrees, and knee fully extended. The footplate was attached to a Systems 2 Dyanomometer (Biodex Medical Systems, Shirley, NY) and Velcro straps were wrapped around the foot to secure it to the plate and prevent movement. Torque was sampled at 2048 Hz and smoothed offline with a 10 Hz low-pass filter (fifth-order Butterworth filter).

#### High-density surface electromyography (HDsEMG)

Before electrode placement, excess hair was removed and the skin overlying the muscle of interest was lightly abraded. High-density surface EMG (HDsEMG) electrodes (64 electrodes, 13×5, 8mm I.E.D., GR08MM1305, OT Bioelettronica, Inc., Turin, IT) were placed over the tibialis anterior (TA), and the medial gastrocnemius (MG) and soleus (SOL), which are dorsiflexor and plantarflexor muscles of the ankle, respectively. A reference electrode strap was placed around the right ankle, overlying the medial and lateral malleolus (Figure 1). HDsEMG data was recorded in single differential mode, sampled at 2048 Hz, amplified x150, and band-pass filtered at 10-500Hz using a Quattrocento signal amplifier (OT Bioelettronica, Inc., Turin, IT). EMG and torque were temporally synced with a 1-second TTL pulse transmitted to the Quattrocento at the onset of each trial.

**Figure 1.**
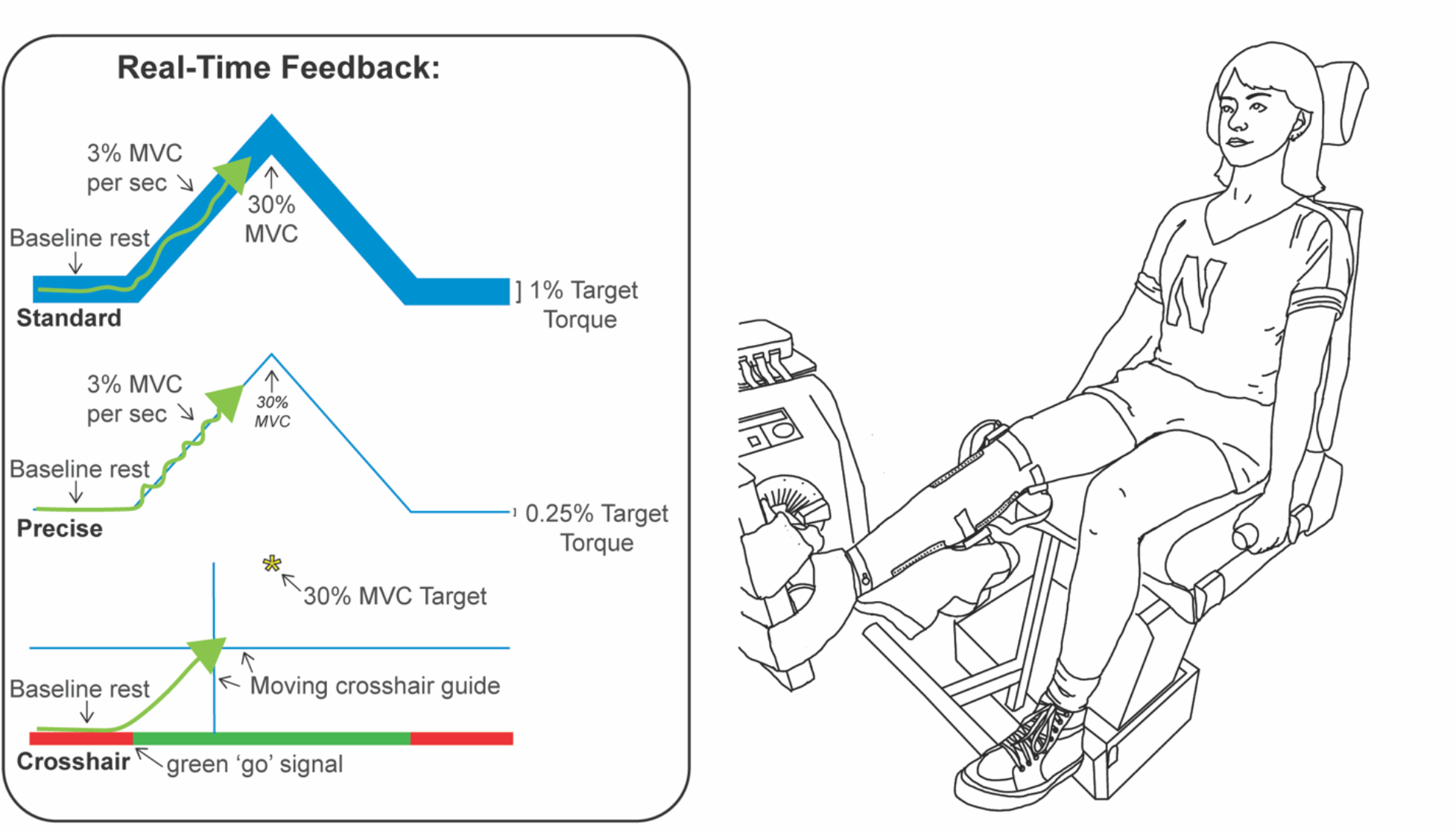
Experimental setup and visual feedback conditions are shown above. Participants were seated with their right ankle secured in a device to measure isometric joint torque, with HDsEMG placed on their tibialis anterior, medial gastrocnemius, and soleus muscles. During the experiment real-time visual feedback was provided on a monitor in front of participants, with three different types of feedback, to cue triangular ramp contractions.

### Experimental Protocol

Before beginning data collection, real-time HDsEMG recordings were visually examined to ensure a high signal-to-noise ratio. First, participants were asked to perform maximum voluntary contractions (MVCs) with their plantarflexors and dorsiflexors. Three MVC trials in each direction were collected. If the last trial had the highest peak torque, additional trials were added as needed. Verbal encouragement was given to participants during MVC trials to ensure maximal performance, and participants were given 2 minutes of rest between trials to prevent fatigue. The maximum torque achieved was used for subsequent normalization of all trials.

The remaining trials consisted of a single triangular-shaped isometric dorsiflexion or plantarflexion torque ramp, with visual feedback to guide participants. For all ramps, participants increased torque (3% MVC/s) to 30% MVC over 10 seconds and then decreased at the same rate to 0% MVC. Throughout the experiment, a large monitor was positioned in front of the participants to provide real-time feedback of torque performance.

#### Visual feedback

This dataset was obtained from a larger ongoing project, and therefore participants performed contractions with three different types of visual feedback, all of which had the same goal of a 10 s linear ascending phase to a peak of 30% MVC and then a 10 s linear descending phase to rest. During all three of these conditions, participants received real-time torque feedback on the screen roughly two meters in front of them, which was comprised of either: 1) a standard guideline with a width of 10% of target torque (i.e., 3% MVC); 2) a precise guideline with a width of 2.5% of target torque (i.e., 0.75% MVC); or 3) cross hairs to provide feedback about the amplitude and timing of torque generation, along with cues about when to start and stop contracting and a peak target (see Figure 2). At a minimum, participants completed 4 trials while receiving each of the three types of feedback in each direction (i.e., plantarflexion and dorsiflexion), yielding 12 ramps for both dorsiflexion and plantarflexion. Trials were grouped by direction and randomized such that participants either performed all plantarflexion first or all dorsiflexion first. Within each direction, the three types of feedback were randomized. Trials were discarded if they did not exhibit a smooth increase and decrease of torque (i.e., substantial deviations from the intended target profile), and the trial was then repeated.

**Figure 2.**
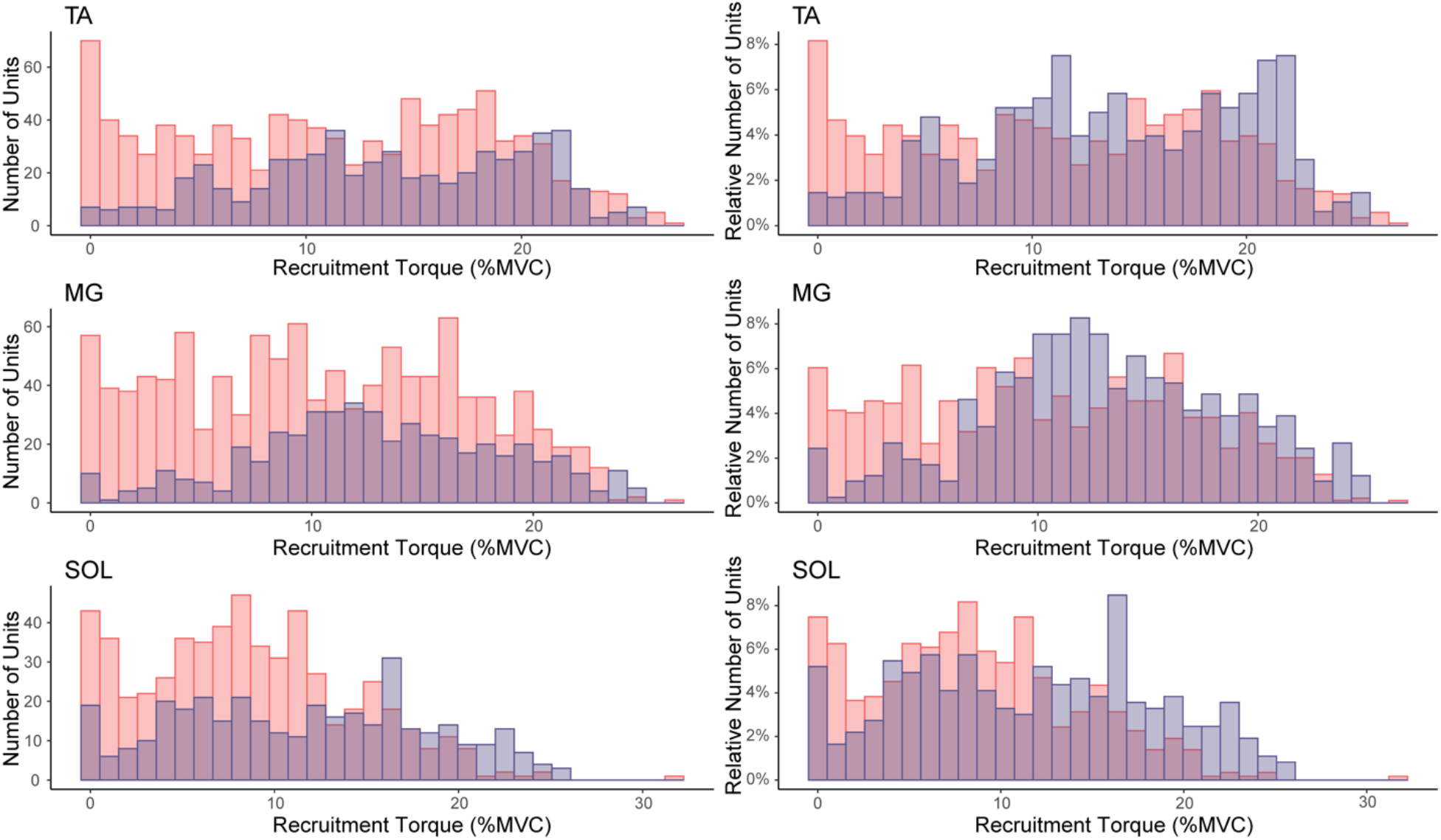
Decomposed MUs by recruitment threshold for all participants for each muscle. Male units are shown in pink, female units overlayed in purple. For each figure, the x-axis shows the percentage of MVC the units were recruited in 1% bins. In the left panels, the y-axis represents the total number of units recruited, whereas in the right panels, the y-axis represents the percentage of total units recruited at each torque level.

### Data Analysis

#### MU Decomposition

After data collection, EMG signals were manually inspected. Channels with substantial artifacts, noise, or analog to digital saturation were removed (on average 1-3 channels per array). Individual MU spike trains were decomposed using a convolutive blind-source separation algorithm with a silhouette threshold of 0.87 (Negro *et al*., 2016).

After decomposition, MU spike trains were manually edited by a trained technician who was blind to the participant’s sex. This inspection used a custom-made graphical user interface in MATLAB to correct minor errors made by the decomposition algorithm using well-validated local re-optimization methods to improve MU spike trains similar to the techniques used in recent studies (Hug et al, 2021; Boccia *et al*., 2019; Afsharipour *et al*., 2020; Del Vecchio *et al*., 2020; Hassan *et al*., 2020; Martinez-Valdes *et al*., 2020). Instantaneous discharge rates of each MU spike train were determined by computing the inverse of the interspike interval and smoothed using support vector regression (Beauchamp *et al*., 2022) with custom-written MATLAB scripts. Within these scripts, the initial, peak, and final discharge rates were extracted from the smoothed spike trains. Ascending duration was calculated as the time that a MU exhibited sustained discharge before peak torque, and descending duration as the time a MU exhibited sustained discharge from peak torque to derecruitment. Ascending-descending phase duration ratio was calculated as the difference in ascending duration and descending duration, divided by the total duration of discharge. A more negative ratio indicates greater duration on the descending portion of the contraction, and thus more sustained discharge on the descending portion of the torque ramp. Finally, recruitment threshold was calculated as the level of torque at the first MU spike.

#### Estimates of Persistent Inward Currents (PICs)

Effects of PICs on MU discharge patterns can be estimated by quantifying the amount of onset-offset hysteresis (i.e., ΔFrequency, ΔF) of a higher-threshold (test) MU with respect to the discharge rate of a lower-threshold (reporter) MU (Gorassini *et al*., 1998, 2002*a*, 2002*b*). Rather than providing ΔF values for each test-reporter unit pair, we calculated ‘unit-wise’ values, which gives each test unit one ΔF value based on the average values obtained from multiple reporter units (Figure 3). Criteria for inclusion of ΔF values from MU pairs were that 1) the test MU was recruited at least 1 second after the reporter unit to ensure full activation of the PIC (Bennett *et al*., 2001; Powers *et al*., 2008; Hassan *et al*., 2020), test unit-reporter unit pair exhibited rate-rate correlations of r^2^ > 0.7 to ensure MU pairs received common synaptic input (Gorassini *et al*., 2004; Udina *et al*., 2010; Stephenson & Maluf, 2011; Vandenberk & Kalmar, 2014), and 3) the reporter unit modulated its discharge rate by at least 0.5 pps while the test unit was active (Stephenson & Maluf, 2011).

**Figure 3.**
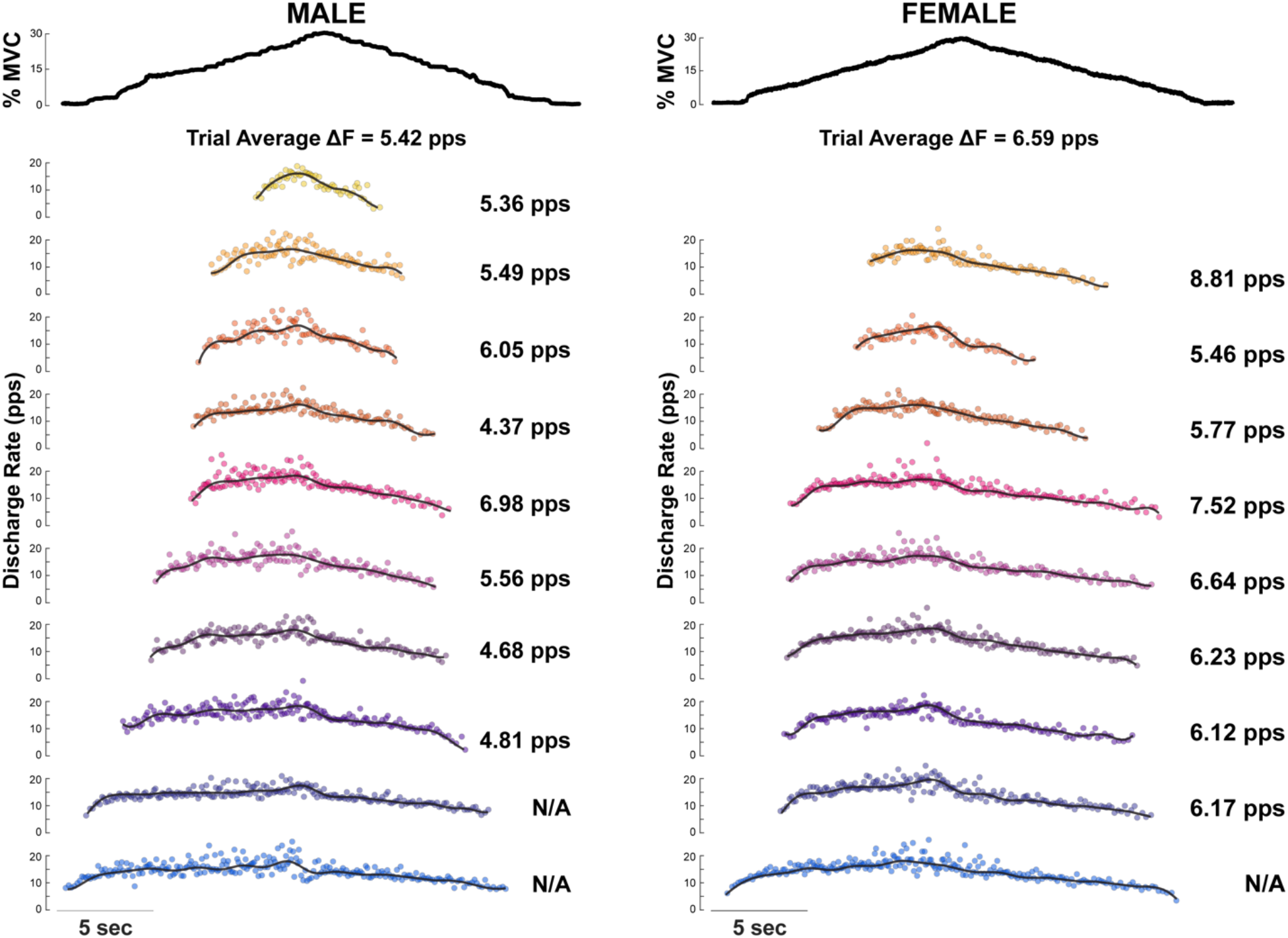
An example of a dorsiflexion contraction trial from one male (left) and one female (right) during which motor units were decomposed from high-density surface electromyography over the tibialis anterior muscle. Torque is plotted with a thick black line (top) and all individual motor units recruited during that trial are plotted in the bottom plots according to recruitment threshold (lower thresholds at the bottom). Each point represents the instantaneous discharge rate, and the thin black line for each unit represents the smoothed discharge rate computed using support vector regression. The test-unit mean, averaged across all suitable reporter units, is provided to the right of each unit. The average of all these values within the trial is provided just below the torque trace.

### Statistical Procedures

Statistical analyses were performed using R Statistical Software (v4.1.0; R Core Team 2021). To determine if variables of interest were predicted by the fixed effects of sex, muscle, and their interactions we used linear mixed effects models with covariates of recruitment threshold and the type of visual feedback, and a random intercept and muscle slope for participant ((Muscle | Participant); lmer R package; v1.1.27.1; Bates *et al*., 2015). To determine significance, we applied Satterthwaite’s method for degrees of freedom (lmerTest R package; v3.1.3; Kuznetsova *et al*., 2017). Both individual participant means and group estimated marginal means (male and female) were computed for each variable (emmeans R package, v1.8.0, Lenth *et al*., 2022). Results are reported as mean ± standard deviation. Effect size (Cohen’s d) was calculated to determine the standardized magnitude of the effect of sex from the estimated marginal means of the male and female data from the model (emmeans R package, v1.8;0, Lenth *et al*. 2022). All data were visualized in R (ggplot R package, v3.3.6, Wickham *et al*., 2022).

#### Comparison between sexes with resampled MUs of matched recruitment torques

The inclusion of recruitment torque as a covariate in the linear mixed effects model indicates that estimates of PICs were higher in females than males across all three muscles despite the limited number of low threshold MUs we decomposed in females. However, to further investigate the possibility that a difference in recruitment threshold between the sexes influenced estimates of PICs and peak discharge rate, we iteratively resampled MUs between the sexes, and compared ΔF and peak discharge rate with similar distributions of recruitment torques. To perform this resampling procedure, for each muscle, we randomly sampled 500 MUs from each sex such that their distributions of recruitment threshold matched each other and represented the mean and standard deviation of MUs across both sexes. To ensure that distributions and collected MUs were not identical with each iteration, we offset the desired mean by a value randomly chosen from a gaussian distribution with a mean of zero and standard deviation of one-quarter of the population standard deviation. Within each iteration, we quantified ΔF and peak discharge rate for each of the MUs in the new distributions and used a linear mixed effects model with a fixed effect of sex and nested random effect of trial number within participant for sex (1 | Sex : PID : Trial Number; fitlme MATLAB package) to determine if sex was a significant predictor of ΔF and peak discharge rate. We repeated this process 1,000 times to determine the probability of significant sex differences, which we report as a value from 0 (0% of runs with a significant difference between sexes) to 1 (100% of runs with a significant difference between sexes).

## RESULTS

In this study, we characterized sex differences in MU discharge properties during isometric triangular shaped contractions to 30% of MVC. For females, we decomposed a total of 2,126 MUs (TA: 933, MG: 691, SOL: 502), and for males a total of 3,935 MUs (TA: 1651, MG: 1484, SOL: 800). Table 1 shows the average number of units decomposed per trial for each muscle and sex.

**Table 1.**
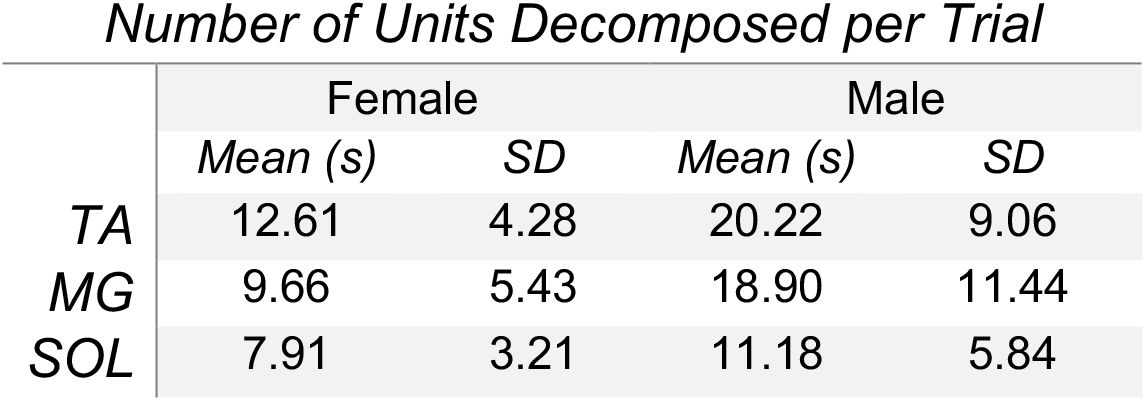
The mean and standard deviation (SD) of units decomposed per trial for each muscle for females and males.

| Number of Units Decomposed per Trial |  |  |  |  |
| --- | --- | --- | --- | --- |
|  | Female |  | Male |  |
|  | Mean (s) | SD | Mean (s) | SD |
| TA | 12.61 | 4.28 | 20.22 | 9.06 |
| MG | 9.66 | 5.43 | 18.90 | 11.44 |
| SOL | 7.91 | 3.21 | 11.18 | 5.84 |

### A greater number of lower threshold MUs are identified in males

When decomposed MUs were stratified by recruitment threshold, it becomes apparent that a greater number of lower threshold MUs were decomposed in males than females (Figure 2). Thus, sex was a significant predictor of recruitment torque (χ^2^_(1)_ = 13.56, *p* = 0.00023), and was higher in females across all three muscles (Figure 2). To account for this in all subsequent analyses, we included a covariate of recruitment torque in our statistical models.

### Estimates of persistent inward currents are greater in females

For both sexes, estimates of PICs were in the normal range of values reported in previous studies (Afsharipour *et al*., 2020. Kim *et al*., 2020). Across all muscles, ΔF was larger (χ^2^_(1)_ = 6.26, *p* = 0.012) in females (4.73 ± 0.242 pps) than males (3.81 ± 0.240 pps) with an effect size of d = 0.653. Figure 3 shows an example dorsiflexion trial from one male and one female, with individual ΔF values provided for the average of all test units and individually for each MU decomposed from the TA muscle. In Figure 4, within participant means (diamonds) and individual ΔF values for each test unit along with distributions across all three muscles are shown. No significant interaction between sex and muscle was identified (χ^2^_(2)_ = 1.79, *p* = 0.409), and therefore ΔF is estimated 0.92 pps higher for all muscles in females based on estimated marginal means.

**Figure 4.**
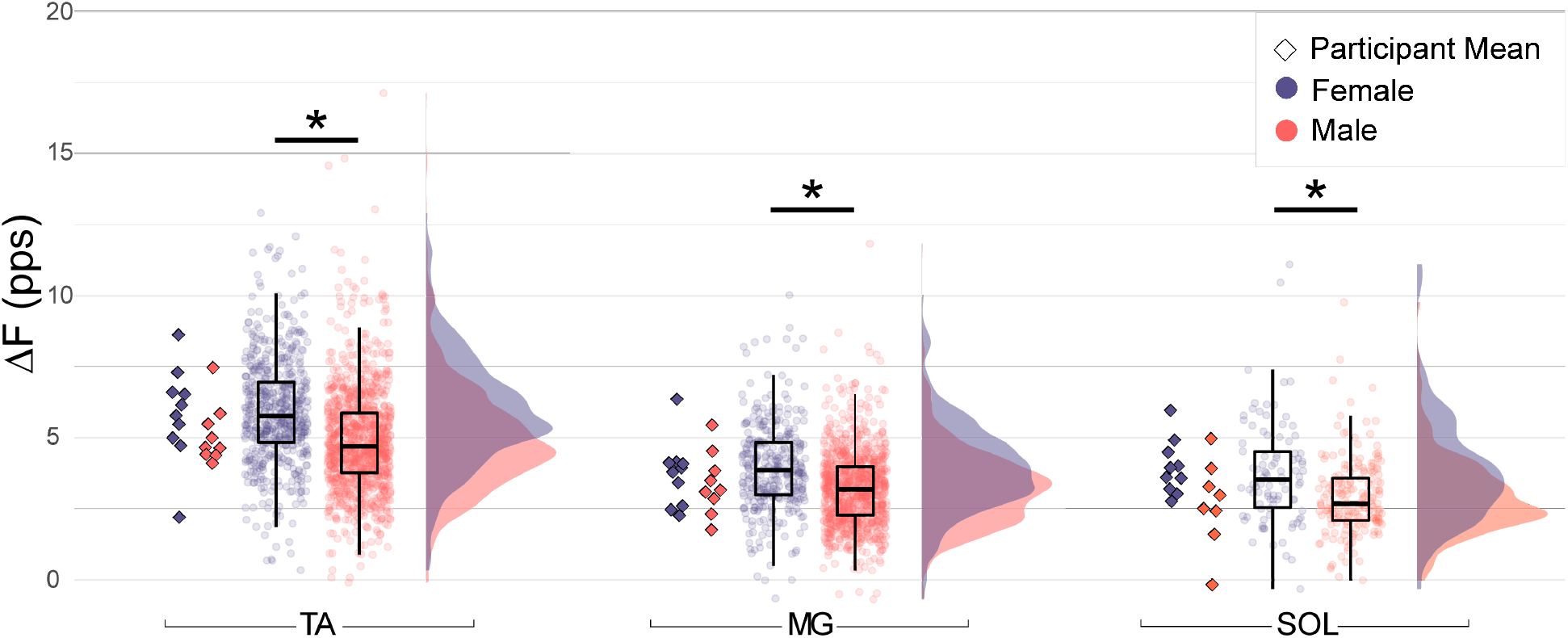
Estimates of persistent inward currents (ΔF) for TA (left), MG (middle), and SOL (right) for females (purple) and males (pink). Individual participant means are shown in diamonds. ΔF measures for each unit collected across trials are shown as scatter points. Box plots represent the 25^th^, 50^th^ (median) and 75^th^ quartiles, with whiskers for minimums and maximums at 1.5 the interquartile range. Distributions of unit-wise ΔF values irrespective of participant are shown to the right for each muscle.

The descending phase duration of discharge, from peak torque to derecruitment (χ^2^_(1)_ = 3.316, *p* = 0.0686), and ascending duration, from recruitment to peak torque (χ^2^_(1)_ = 0.0342, *p* = 0.8532), were similar between the sexes (Table 2B). Additionally, the ratio between ascending and descending durations did not differ significantly between the sexes (χ^2^_(1)_ = 2.94, *p* = 0.086, Table 2C).

**Table 2.** Measurements of motor unit discharge duration during the triangular ramp contractions. Estimated marginal means and standard deviations are shown for each muscle, with females on the left and males on the right. Abbreviations: TA; tibialis anterior, MG; medial gastrocnemius, SOL; soleus, s; seconds, SD; standard deviation.

A. Ascending Phase Duration
|  | Female |  | Male |  |
| --- | --- | --- | --- | --- |
|  | Mean (s) | SD | Mean (s) | SD |
| TA | 6.09 | 0.52 | 6.11 | 0.52 |
| MG | 5.97 | 0.51 | 5.98 | 0.51 |
| SOL | 6.03 | 0.51 | 6.09 | 0.52 |

B. Descending Phase Duration
|  | Female |  | Male |  |
| --- | --- | --- | --- | --- |
|  | Mean (s) | SD | Mean (s) | SD |
| TA | 7.32 | 1.72 | 6.58 | 1.74 |
| MG | 5.23 | 1.88 | 4.48 | 1.87 |
| SOL | 5.77 | 1.78 | 5.02 | 1.85 |

C. Ascending-Descending Phase Ratio
|  | Female |  | Male |  |
| --- | --- | --- | --- | --- |
|  | Mean (s) | SD | Mean (s) | SD |
| TA | -0.16 | 0.20 | -0.02 | 0.20 |
| MG | 0.12 | 0.25 | 0.12 | 0.25 |
| SOL | 0.0369 | 0.22 | 0.0817 | 0.24 |

### MU Discharge rates did not differ between sexes

While peak discharge rates appeared higher in females than males (see table 3A), no significant differences were detected between the two groups. Across muscles, discharge rates did not differ at recruitment (χ^2^_(1)_ = 1.36, *p* = 0.243), derecruitment (χ^2^_(1)_ = 0.06, *p* = 0.805), or at the peak (χ^2^_(1)_ = 1.15, *p* = 0.283).

**Table 3.** Motor unit discharge rates during triangular ramp contractions. Estimated marginal means and standard deviations are shown for each muscle, with females on the left and males on the right. Abbreviations: TA; tibialis anterior, MG; medial gastrocnemius, SOL; soleus, pps; pulses per second; s; seconds, SD; standard deviation.

A. Discharge Rate at Recruitment
|  | Female |  | Male |  |
| --- | --- | --- | --- | --- |
|  | Mean (pps) | SD | Mean (pps) | SD |
| TA | 6.73 | 1.98 | 6.29 | 2.01 |
| MG | 5.01 | 1.91 | 4.57 | 1.96 |
| SOL | 4.04 | 1.83 | 3.60 | 1.88 |

B. Peak Discharge Rate
|  | Female |  | Male |  |
| --- | --- | --- | --- | --- |
|  | Mean (pps) | SD | Mean (pps) | SD |
| TA | 17.20 | 2.33 | 16.40 | 2.41 |
| MG | 13.00 | 2.72 | 12.10 | 2.76 |
| SOL | 11.10 | 2.43 | 10.30 | 2.57 |

|  | Female |  | Male |  |
| --- | --- | --- | --- | --- |
|  | Mean (pps) | SD | Mean (pps) | SD |
| TA | 5.55 | 1.61 | 5.50 | 1.62 |
| MG | 4.60 | 1.63 | 4.56 | 1.64 |
| SOL | 3.70 | 1.58 | 3.65 | 1.60 |

### Resampling recruitment threshold distributions

To account for the differences in recruitment thresholds, we resampled 1000 iterations of random MUs with a similar distribution of recruitment threshold between males and females showed that ΔF was still significantly higher in females than males (Fig. 5B-C). Across the 1000 iterations, we found a probability of obtaining significant differences in ΔF of 0.98, 1.00, and 1.00 for the MG, SOL, and TA, respectively. Furthermore, we found the average beta coefficients for the fixed factor of sex across all runs to be 0.63 ± 0.08, 1.01 ± 0.1, and 1.06 ± 0.1 for the MG, SOL, and TA, respectively. Resampling also revealed that peak discharge rates were higher in females for MG and SOL, but not TA (Fig. 5D-E), with the probability of significant differences of 0.63, 1.00, and 0.00, respectively. The fact that the resampled data with similar recruitment thresholds revealed higher discharge rates in females likely reflects the larger numbers of low threshold units in males compared to females (see Discussion).

**Figure 5.**
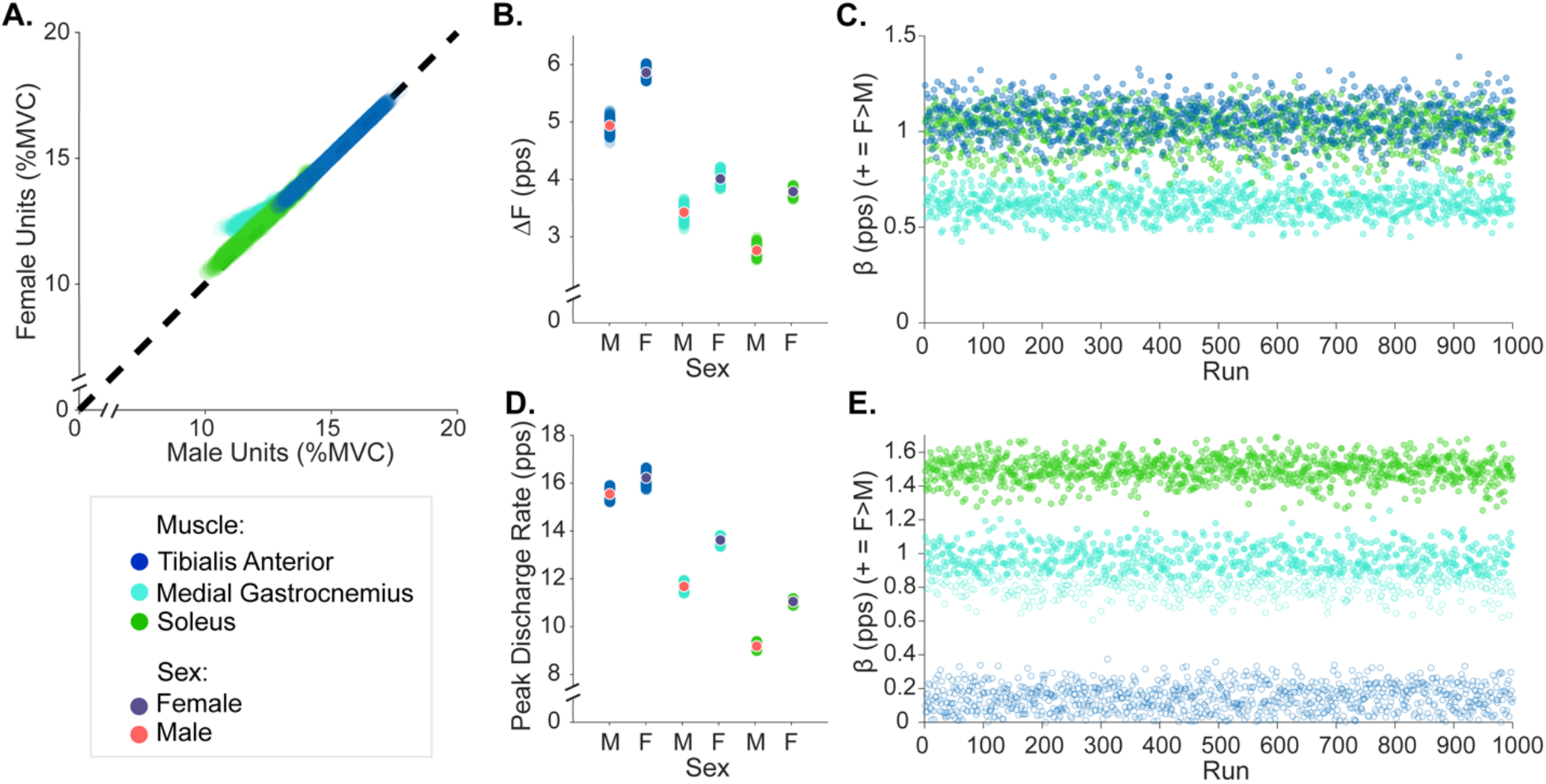
Estimates of persistent inward currents (ΔF) and peak discharge rate between females (F) and males (M) based on resampled MUs with equal distributions of recruitment threshold between the sexes. **A**. Average recruitment threshold, percent maximum voluntary contraction (% MVC), of resampled MUs for each sex across 1,000 runs for each muscle along the line of identity. **B**. Unit-wise and average ΔF values from resampled MUs for each muscle and sex. **C**. Beta coefficients (β), indicating the difference between female and male ΔF for each run, with filled points representing significant values and unfilled indicating non-significant values. **D**. Individual and average peak discharge rate from resampled MUs for each muscle and sex. **E**. Beta coefficients (β), indicating the difference between female and male peak discharge rates for each run, with filled points representing significant values and unfilled indicating non-significant values.

## DISCUSSION

This study quantified the discharge patterns of several concurrently active MUs during submaximal isometric triangular-shaped contractions in the lower limb muscles of females and males. Although no significant differences in MU discharge rates were identified, estimates of PIC magnitude using a paired motor unit analysis technique revealed that discharge rate hysteresis (i.e., ΔF) was significantly larger in females than males. These findings suggest that descending neuromodulatory drive, monoaminergic signaling of 5HT and/or NE to motor units, and/or inhibitory inputs may differ between the sexes.

### More lower threshold MUs decomposed in males than in females

In the current investigation, we identified a greater number of MUs in males across all three of the sampled muscles. Recent work by our colleagues (Taylor *et al*., 2022) also documented a higher number of Mus decomposed from the TA muscle in males compared to females, and our findings extend these findings to other muscles of the lower leg. In particular, a greater proportion of lower threshold MUs were decomposed in males compared to females in the current study.

Methodological considerations that may skew comparisons of recruitment thresholds between heterogeneous groups of people (i.e., females vs males herein) should be fully considered before in-depth interpretations are made. For example, HDsEMG decomposition is biased toward larger and more superficial MUs (Hassan *et al*., 2019) and, as shown by Guo and colleagues (Guo *et al*., 2022), males tend to have larger MUs, which may explain why their spike times are decomposed more readily than their female counterparts (Taylor *et al*., 2022). These sex differences in the decomposition results are important to note because they highlight a barrier to female inclusion in past HDsEMG analyses. To account for differences in recruitment threshold on ΔF and peak discharge rate, we ran a model to resample MUs with similar distributions of recruitment torque to compare these measures between the sexes. Both the comparisons of our entire dataset and the resampled data with comparable recruitment thresholds are considered in our interpretations and discussion below.

### Estimates of PICs are higher in females

Linear mixed effects models revealed that biological sex was a significant predictor of estimates of PIC magnitude (ΔF) across all three muscles. Indeed, ΔF was almost 1 pps higher in females than males, indicating a greater contribution of PICs to MU discharge while performing a simple isometric ramp contraction. Resampling of MUs to compare estimates of PICs with equal distributions of recruitment torque agreed with this finding, confirming that less low threshold MUs decomposed in females did not affect estimates of PICs and comparisons between the sexes.

PICs are vital for motoneuronal function – without their amplification effects, repetitive discharge cannot be achieved (Lee & Heckman, 2001), as driving currents from descending and sensory synaptic inputs are too weak to cause repetitive MU discharge (Binder *et al*., 2020). Therefore, identifying and understanding differences in PIC magnitude may garner insights into potential sex differences in motor command structure. Even though males in our experiment certainly displayed discharge rate hysteresis (i.e., positive ΔF values), the magnitude of this hysteresis was markedly smaller than females.

The factors that can influence PIC magnitude, and thus ΔF, include: 1) monoaminergic input to the motoneurons, 2) Na+ or Ca2+ channel function, and/or changes in 5HT/NE reuptake and receptor function, and 3) the amount or pattern of inhibitory input to the motoneuron. Physiologically, males and females differ most notably in their levels of sex hormones. In eumenorrheic females (premenopausal females with a regular menstrual cycle), levels of estradiol, one of the three endogenous estrogens, and progesterone fluctuate throughout the month over a menstrual cycle (Sherman & Korenman, 1975). In males, the levels of these two hormones remain low and do not fluctuate across the month. Progesterone levels in males are similar to levels of females during the early menstrual cycle (follicular phase), but after ovulation, progesterone levels rise much higher in females (Oettel & Mukhopadhyay, 2004; Tea *et al*., 1975). Throughout the entire menstrual cycle, eumenorrheic females experience fluctuations in estradiol ten times that of males (Verdonk *et al*., 2019). Remarkably, these two hormones have known effects on neurons throughout the nervous system, in particular serotonin (Bethea *et al*., 1998), and thus, higher fluctuating levels in females may have effects on the motor system.

### MU discharge rates did not differ between males and females

Despite the tendency for females to have higher discharge rates overall, there were no significant sex differences in MU discharge rates when considering our entire dataset. Previous studies have reported higher discharge rates in female MUs, however, during these experiments participants performed isometric holds or trapezoid-shaped contractions (Inglis & Gabriel, 2020; Guo *et al*., 2022; Taylor *et al*., 2022). In our study, discharge rates at onset, maximum, and offset did not differ between the sexes. The greater yield of lower threshold units in the males may have skewed the discharge rates of males to a higher level, due to the relationship of recruitment threshold and discharge rate during submaximal isometric contractions (Pearcey & Rymer, 2022), which would diminish statistical differences between the sexes. We did, however, attempt to deal with this limitation by including a covariate of recruitment threshold in our models. It remains possible that, if participants had been asked to sustain a contraction, sex differences may have emerged as has been observed by previous studies. Moving one step beyond a covariate in our analysis, the peak discharge rate in our resampled data with equal distributions of recruitment torque revealed muscle specific differences between the sexes. The resampling of MUs revealed that females had greater peak discharge rate in the MG and SOL, suggesting that peak discharge rates may be higher in females when matched for recruitment torque in the plantarflexors, not in the dorsiflexors (i.e., TA).

### The potential role of sex hormones in mediating PIC magnitude

A possible underlying mechanism that could explain higher PICs in females is the interaction between monoamines released from the caudal raphe nucleus (i.e., 5HT) and locus coeruleus (i.e., NE) and the sex hormone estradiol. These monoamines facilitate PICs, and therefore the magnitude of PICs is directly proportional to the level of brainstem monoamine release (Lee & Heckman, 1998, 2000). Thus, sex differences in monoaminergic release could be responsible for differences in ΔF, or perhaps the effects of estradiol on 5HT neurons in other areas of the nervous system. In nonhuman female primates, estrogen receptor beta (Erβ) has been identified in the dorsal and caudal raphe nuclei (Gundlah *et al*., 2000; Bethea *et al*., 2002). Long-term loss of estradiol in ovariectomized female macaques also results in a reduction in the number of 5HT neurons (Rivera & Bethea, 2012). Taken together, these data suggest that females could have greater release of brainstem-derived monoamines than males due to greater estradiol effects in the brainstem.

The hormones estradiol (one of the three endogenous estrogens) and progesterone fluctuate across the female menstrual cycle in young adult females, and peak at levels much higher than males during specific days (Sisk & Foster, 2004). Thus, since this work has revealed potential sex differences in monoaminergic signaling, an essential next step will be to determine the effect of fluctuating levels of estradiol and progesterone on estimates of PICs. If sex differences were in fact caused by differences in sex hormones, and since we did not control or measure the time of the menstrual cycle in females, we would expect greater variability in estimates of PICs across the female group due to the fluctuating levels of sex hormones in females in comparison to the relatively stable levels in males. Additional examination of values obtained across individuals within each sex indeed suggests that there is more variability in ΔF and discharge rates in the group of females compared to males. Measures of variance in ΔF obtained from our mixed-effects models were indeed greater in females (see Figure 4). Confidence in this sentiment, however, requires further study, in which females are studied at multiple time points throughout their menstrual cycle, and sex hormones are quantified.

### Methodological considerations

The non-invasive nature of surface electromyography is advantageous in that it allows us to estimate the spike times of many MUs but since we are unable to measure from the spinal motoneurons directly, this method is unable to directly quantify the magnitude of PICs. The paired MU analysis technique to estimate PIC magnitude in humans has been well-validated, but it is possible that the mechanisms behind the observed sex differences in ΔF are not solely due to changes in monoaminergic drive. For example, differences in the patterns of inhibitory inputs, monoamine release, monoaminergic receptor activity and/or sensitivity differences between the sexes could possibly account for differences in MU discharge patterns (Beauchamp *et al*., 2023). Future studies which include a comprehensive array of contractions, that complement the isometric triangular ramps utilized herein, and allow inferences about the biophysical properties of motoneurons (Khurram *et al*., 2022) or that use electrical stimulation or vibration protocols to probe excitatory and inhibitory circuits (Mesquita *et al*., 2021, Pearcey *et al*., 2022, Orssatto *et al*., 2022), may reveal further insight into the mechanism behind these sex differences. A complimentary approach that also incorporates measures of musculoskeletal properties and/or mechanical/passive properties of the muscle during the contraction (Martinez-Valdes *et al*., 2022) may further elucidate the mechanisms responsible for sex differences in discharge patterns quantified here (Heroux *et al*. 2020).

Another limitation is that we assessed only one submaximal level of contraction (i.e., 30% MVC). It is possible that sex differences in PICs and discharge rates are dependent on contraction intensity. For example, previous work has shown discharge rates during plateau holds are higher in females until 40% MVC, equal between the sexes at 60 and 80%, and higher in males at 100% MVC (Inglis and Gabriel 2020). When data is strength matched, however, discharge rates in females were higher across all intensities. In the current experiment, estimates of PICs were higher in females during low intensity contractions but it is possible that increasing the contraction intensity could reveal further insight into monoaminergic signaling between the sexes.

Another limitation of this study is that the menstrual cycle status of female participants was not recorded and thus could not be taken into consideration during analysis. The uncontrolled phase of the menstrual cycle in females remains a potential reason for the increased variability in female ΔF values. Previous work has shown that various aspects of motor control, including MVC (Sarwar *et al*., 1996; Nicolay *et al*., 2007), fatigability (Sarwar *et al*., 1996; Ansdell *et al*., 2019; Pereira *et al*., 2020), and initial MU discharge rate (Tenan *et al*., 2013) all fluctuate across the menstrual cycle. There is reason to believe that MU discharge and PIC magnitude would also be affected by fluctuating levels of sex hormones, and thus this study provides motivation to examine these MU properties across the female menstrual cycle.

Subcutaneous fat, or the distance between the muscle and the surface array, is a likely contributor to the lower MU yield in females, however we cannot confirm this because we did not perform appropriate measures for each participant. Larger distances between the electrodes and the muscle fibers in the females recruited for this study (i.e., higher average levels of subcutaneous fat) are likely, based on typical (i.e., population) anthropometrics (Deurenberg *et al*., 1991). However, future analyses that include matching subcutaneous fat thickness between the sexes will offer further insight.

## Conclusion

We have revealed that estimates of PIC magnitudes are higher in the lower limb muscles of females than males. This suggests female MUs receive a greater contribution from PICs than males to achieve the same relative force and discharge rate, which may be due to a requirement for more persistent discharge or a consequence of more monoamine availability. Females may also use a different combination of synaptic excitation, inhibition, and neuromodulation than males to achieve the same relative force levels during triangular ramp contractions. As such, this study further highlights the necessity to include equal numbers of female participants in order to, not only, fully understand the human physiology behind motor control, but to potentially use MU analysis as biomarkers for neurological disease (Fajardo *et al*., 2022).

## Acknowledgements

The authors would like to thank Obaid Khurram for his fruitful discussion during planning of the experiments and Kirsten Carroll for assistance with data analysis.

## Data availability statement

The data that support the findings of this study are available on request from the corresponding author.

## Author contributions

S.T.J., J.A.B., C.J.H., and G.E.P.P., conceptualized and designed the research; S.T.J., J.A.B., M.M.G., and G.E.P.P. performed the experiments; S.T.J. and J.A.B. analyzed the data; S.T.J., J.A.B., G.E.P.P., and C.J.H. interpreted the results of experiments; S.T.J., J.A.B., and G.E.P.P., prepared the figures; S.T.J. drafted the manuscript; S.T.J., J.A.B., M.M.G., G.E.P.P., F.N., and C.J.H. revised and approved the final version of the manuscript.

